# Construction of transcript regulation mechanism prediction models based on binding motif environment of transcription factor AoXlnR in *Aspergillus oryzae*

**DOI:** 10.1101/2021.07.28.454268

**Authors:** Hiroya Oka, Takaaki Kojima, Ryuji Kato, Kunio Ihara, Hideo Nakano

## Abstract

Recent study revealed that there are thousands of genes that remain unaffected by increased AoXlnR expression, despite the presence of one or more AoXlnR-binding motifs in their promoter region. Given this knowledge, we designed this study to construct several predictive models for determining whether a gene can exhibit a differential response to changes in AoXlnR expression. These models were constructed using 3D DNA shape information determined using the sequence around the AoXlnR binding motifs with classification as functional or nonfunctional. These models were created using a support vector machine followed by the evaluations designed to determine whether these DNA shape-based models can correctly classify functional motifs in terms of area under the receiver operating characteristic curve. The results showed that the differential expression levels of genes located downstream of the AoXlnR motif are closely related to specific DNA shape information around the binding motifs. Furthermore, we found that the parameters contributing to differential expressions differed depending on the number of motifs in the promoter region by comparing the prediction models using regions with only one binding DNA motif and those with multiple binding DNA motifs.

**Author Summary:** DNA-binding transcription factors (TFs) play a central role in transcriptional regulation mechanisms, mainly through their specific binding to target sites on the genome and regulation of the expression of downstream genes. Therefore, a comprehensive analysis of the function of these TFs will lead to the understanding of various biological mechanisms. However, the functions of TFs *in vivo* are diverse and complicated, and the identified binding sites on the genome are not necessarily involved in the regulation of downstream gene expression. In this study, we investigated whether DNA structural information around the binding site of transcription factors can be used to predict the involvement of the binding site in the regulation of the expression of genes located downstream of the binding site. Specifically, we calculated the structural parameters based on the DNA shape around the DNA binding motif located upstream of the gene whose expression is directly regulated by the transcription factor AoXlnR from *Aspergillus oryzae*, and showed that the presence or absence of expression regulation can be predicted from the sequence information with high accuracy by machine learning incorporating these parameters.

## Introduction

Transcription factors (TFs) play a central role in complicated and diverse transcriptional regulatory mechanisms [1], therefore, the comprehensive analysis of the functions of TFs is critical to our understanding of various biological mechanisms. TF-DNA interactions have been analyzed by SELEX-Seq [2, 3], ChIP-Seq [4], whereas the effect of TFs on the expression of genes has been mostly evaluated by RNA-Seq and microarray. In order to comprehensively understand complicated regulatory mechanisms mediated by TFs, it is necessary to integrate several biological information acquired by these different techniques for comprehensive visualization of transcriptional regulatory mechanisms [5–7].

The binding preference of a TF is generally expressed as a DNA motif with 10 to 20 bp in length allowing a variety of sequence substitutions. The binding motifs can be analyzed from the collected TF-binding sequences using the position weight matrix (PWM) that calculates the probability of a base appearing at one sequential position independently, followed by the visualization of the probability as a motif logo. However, PWM does not consider interactions between adjacent sequences [8, 9]. Additionally, several studies have shown that the number of TF binding sites intrinsically related to the expression of these regulated genes in cells is significantly less than that of the binding DNA motifs in the genome [10, 11], which suggests the need for developing an alternative, more multifaceted, approach to understand the complex relationship between TFs and their binding sites.

Olson and co-workers first analyzed the crystal structures of complex of various 10 bp DNA double strand and a TF, and determined DNA shape parameters from the calculated average structurers of the DNA-TF complexes [12]. Later, four parameters, HelT, MGW, ProT, and Roll were developed as DNA shape features by Zhou and co-workers using a pentamer-based modelling from all-atom Monte Carlo simulations of DNA structures [13]. A recent study using these parameters showed that recognition sequences of more than 100 TFs were determined by the DNA shape information more accurately than by conventional sequence-based methods, indicating that PWMs ignore the conformational aspects of DNA in TF binding [14]. Additionally, several studies have reported that the inclusion of the DNA shape information enhanced the accuracy of TF-binding prediction models constructed only from DNA sequence information [15, 16]. For example, Dror *et al*. showed that DNA conformational preferences differ between TF families by evaluating DNA shape information around TF-binding sites [17]. However, to date, these studies have primarily focused on only the difference among TF family units, such as C2H2 zinc finger TFs, not on an individual TF. Additionally, no in-depth analysis linking TF preference, DNA shape and regulatory gene expression has yet been reported for most individual TFs, except for one paper [7].

In our previous study, we used integrated experimental data obtained from genomic SELEX-Seq (gSELEX-Seq) and microarray analysis [5, 6] to evaluate all genes transcriptionally regulated by the *Aspergillus oryzae* TF AoXlnR. *In vitro* AoXlnR binding sites were detected in the promoter regions of more than 2000 genes, however, less than 3% of these genes responded to high expression of AoXlnR in *A. oryzae* cells [6]. This result strongly indicates the existence of an unknown factor regulating TF-DNA interaction in addition to the targeted DNA sequences.

In this study, by using gSELEX-Seq and DNA micro array data of AoXlnR used in a previous study [6], we constructed predictive models with a support vector machine to investigate the relationship between the conformations of DNA around the TF-binding motifs for AoXlnR and the expression of downstream genes (Fig. 1). The results of these evaluations showed that AoXlnR-regulated differential expression could be predicted with high accuracy using the DNA shape parameters of the sequence surrounding TF-binding motifs. Furthermore, it was confirmed that the optimal model for predicting the differential expression depends on the number of binding motifs in the promoter region.

**Figure 1.**
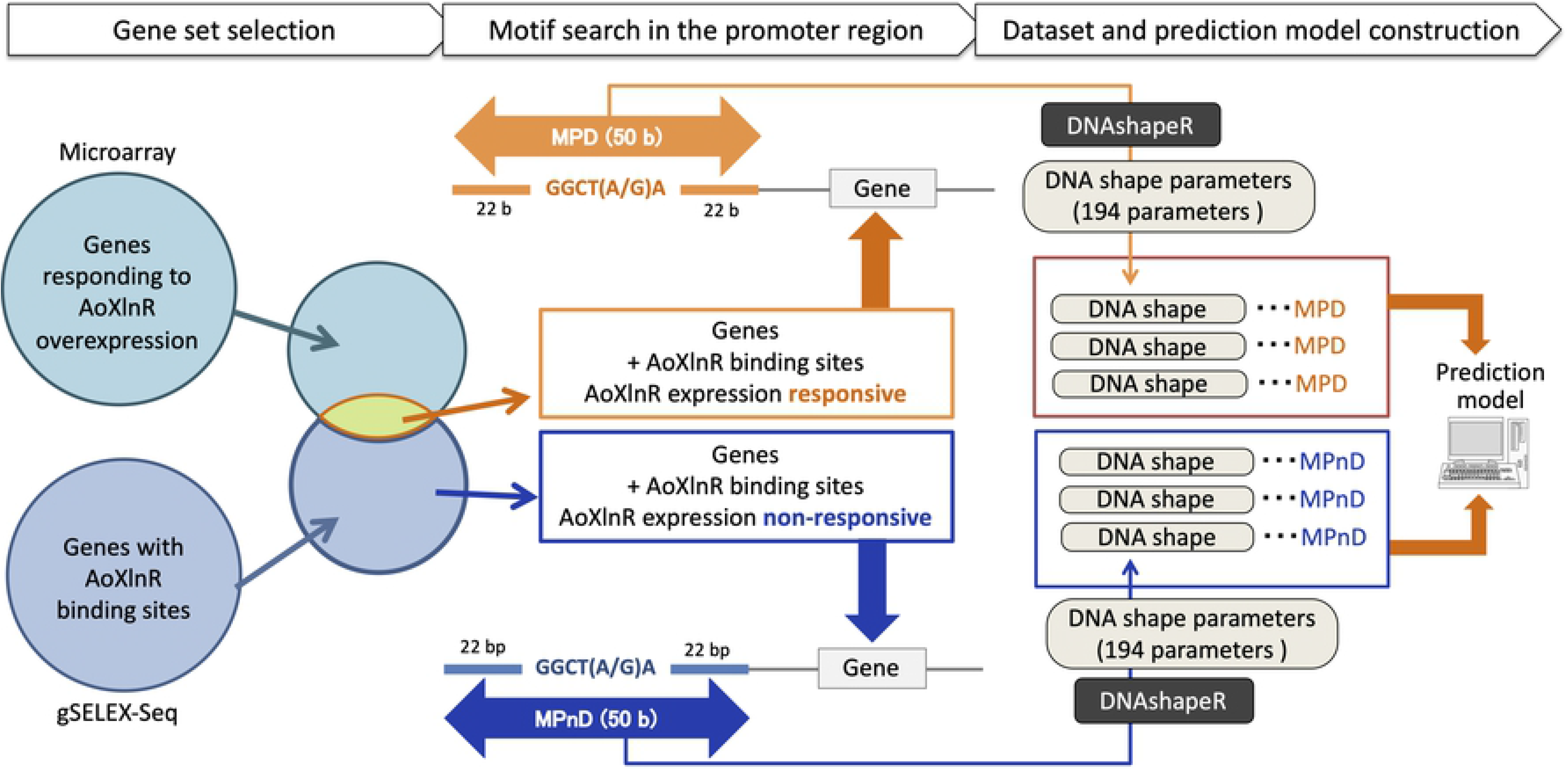
Flowchart describing the construction of the supervised machine leaning models to predict AoXlnR-dependent differential expression from DNA shape parameters. The genes identified in previous studies were classified into two groups depending on the detection method used in their isolation. The positions of AoXlnR binding motifs, 5′-GGCTGA-3′ or 5′-GGCTAA-3′ in each promoter of genes were investigated. Once identified we extracted 22 b up and downstream of these recognition sites and used these to identify the relevant DNA shape parameters. These parameters were then used to construct the discriminative model used in the machine learning programs.

## Results

### Construction of AoXlnR binding motif classification models using DNA shape parameters

We identified 51 genes likely to be directly regulated by AoXlnR by integrating information of the binding sites in *A. oryzae* genome obtained from gSELEX-Seq [6] and the differential expression data obtained from the microarray completed in a previous study [18]. The 194 DNA shape parameters (MGW:48, ProT:48, HelT:49, and Roll:49) were used as explanatory valuables for constructing classifying AoXlnR binding motifs as MPD (63 motifs) or MPnD (1506 motifs). Model performances were evaluated with their AUC values (Fig. 2, Supplementary Fig 1). AUC for the model constructed with selected DNA shape parameters was higher than those for the models that used all the parameters. The selection of contributed DNA shape parameters improved the performance of the prediction. When the same procedure was applied to construct models using the antisense sequences of the AoXlnR binding motifs, the obtained performances using selected shape features were significantly higher than that using all shape features was the case with the original AoXlnR-binding motif sequences (Fig. 2, Supplementary Fig. 1).

**Figure 2.**
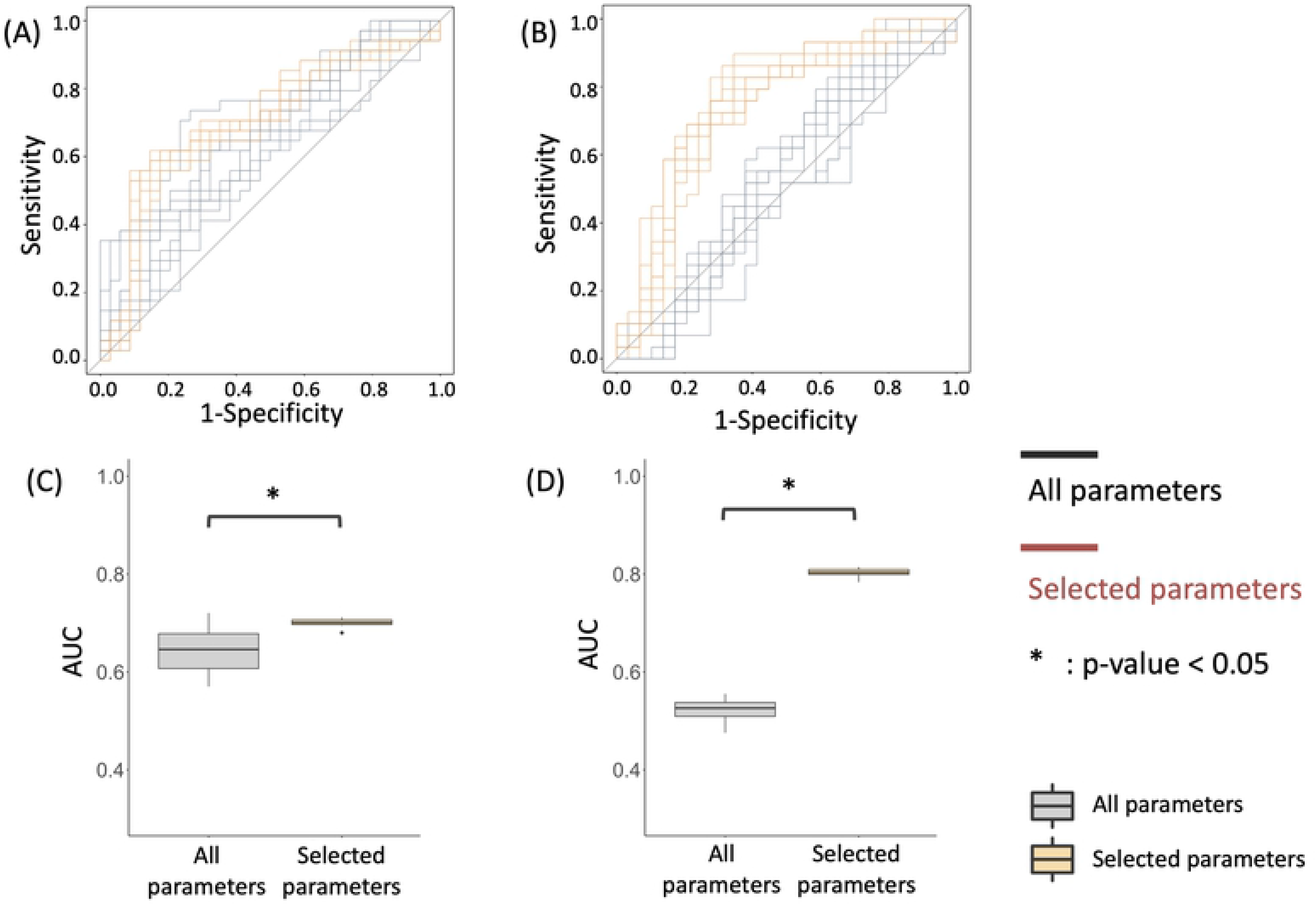
ROC curves and box plots amongst the various models constructed using DNA shape parameters. (A) ROC curve of the model based on the sense strands of MPDs.(B) ROC curve of the model based on the antisense strands of MPDs.(C) Box plots of AUC values obtained from ROC curve of the model based on the sense strands of MPDs.(D) Box plots of AUC values obtained from ROC curve of the model based on the antisense strands of MPDs. Each model was constructed using DNA shape parameters calculated from 50 b sequences containing AoXlnR binding motifs. Here we displayed the ROC curves of the models with the highest AUC for discriminant model using non-stratified MPDs selected parameters shown in Supplementary Figure 1A (for the sense sequences) or 1B (for the antisense sequences). The black lines show the ROCs of the model using all DNA shape parameters, and the red lines show the ROCs of the model using only the preferentially selected parameters shown in Supplementary Figure 1A or 1B. Ten lines show the result of 10 times validation pairing the positive and negative samples in LOOCV. The vertical axis of the ROC curve indicates the “Sensitivity”, which is the true positive rate, and the horizontal axis indicates the “1 - Specificity”, which is the false positive rate. *p* values were calculated using the Student’s *t* test. \**p* < 0.05.

### Stratification of differential expression prediction models based on the number of binding motifs in promoters

The number of recognition sequences in a promoter was suggested to be associated with its responsiveness to that TF previously [6]. Therefore, we stratified MPDs according to the number of AoXlnR motifs present in their promoter regions, separating them into two groups: one motif and two or more motifs.

The DNA shape parameters of the classified datasets were calculated and then used to produce the prediction models as described above. When these models were compared, we found that the models constructed using the top parameters with the highest contribution to the expression differences showed significantly higher AUC values than the models constructed using all the parameters, as in the case of using the non-stratified MPDs (Fig. 3, Supplementary Fig. 2). Additionally, the model constructed using the stratified dataset showed higher AUC values than those constructed using the unclassified datasets. In particular, the classification by the number of motifs tended to more highly contribute to increased AUC values when analyzing the antisense sequences of the AoXlnR binding motifs (Figs. 3B, D, F and H, Supplementary Figs. 2B and 2D).

**Figure 3.**
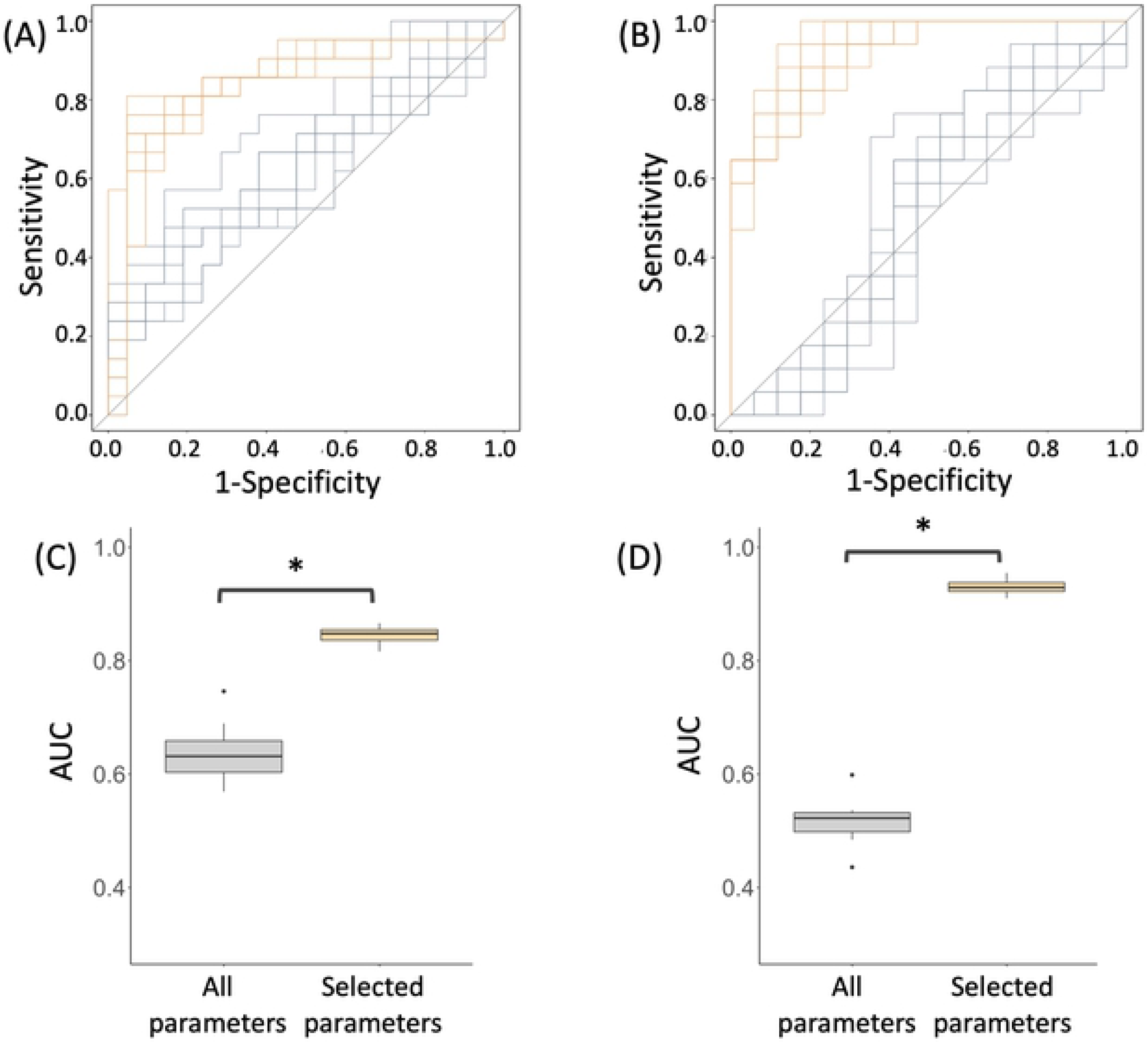
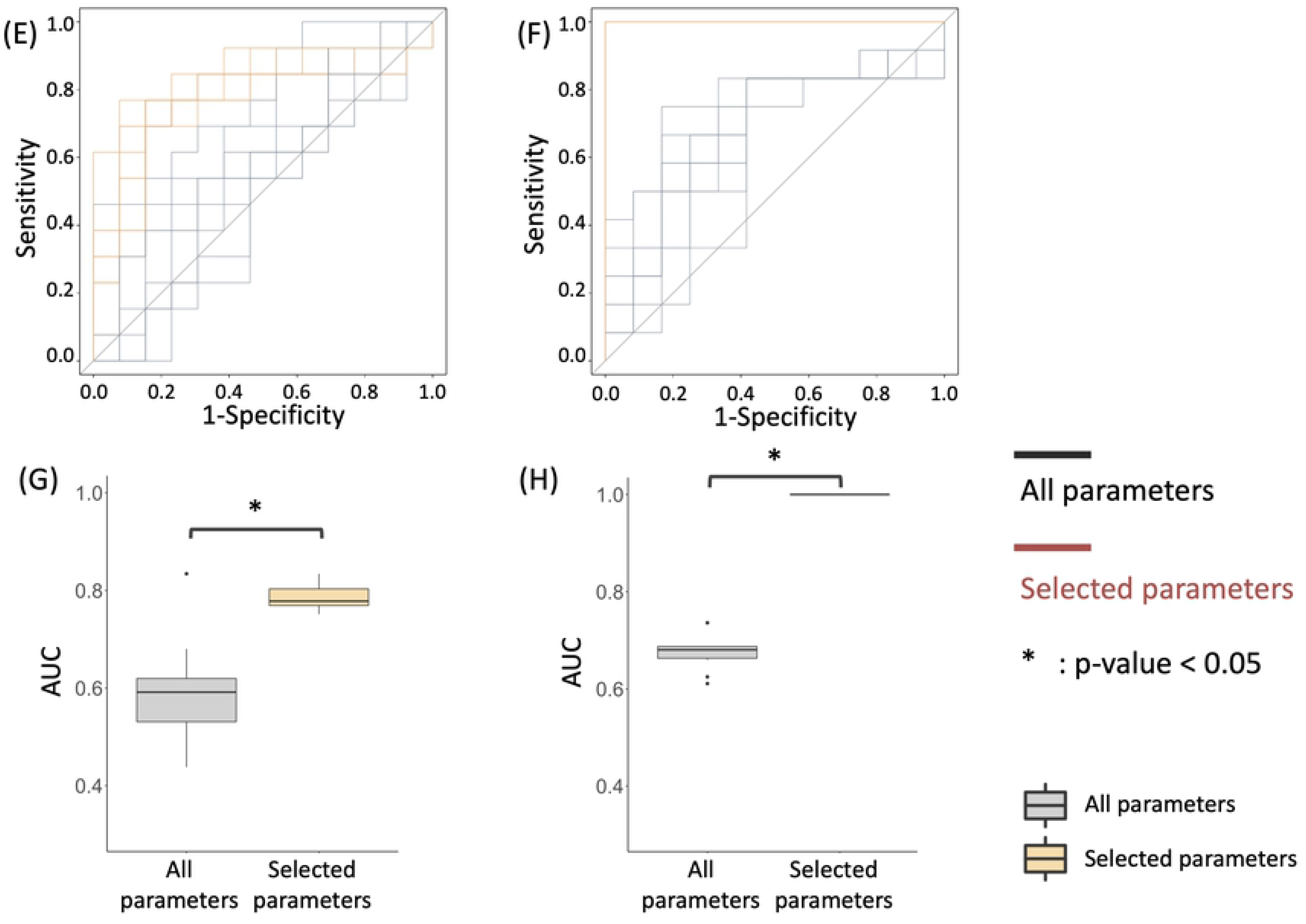
ROC curves and box plots amongst the various models constructed using stratified dataset of DNA shape parameters. (A) ROC curve of the model based on the sense strands of MPDs from the promoters including a single motif.(B) ROC curve of the model based on the antisense strands of MPDs from the promoters including a single motif.(C) Box plots of AUC values obtained from ROC curve of the model based on the sense strands of MPDs from the promoters including a single motif.(D) Box plots of AUC values obtained from ROC curve of the model based on the antisense strands of MPDs from the promoters including a single motif.(E) ROC curve of the model based on the sense strands of MPDs from the promoters including multiple motifs.(F) ROC curve of the model based on the antisense strands of MPDs from the promoters including multiple motifs.(G) Box plots of AUC values obtained from ROC curve of the model based on the sense strands of MPDs from the promoters including multiple motifs.(H) Box plots of AUC values obtained from ROC curve of the model based on the antisense strands of MPDs from the promoters including multiple motifs. Each model was constructed using DNA shape parameters calculated from 50 b sequences containing AoXlnR binding motifs. Here we displayed the ROC curves of the models with the highest AUC for discriminant model using stratified MPDs selected parameters shown in Supplementary Figure 2A (for the sense sequences including a single motif), 2B (for the antisense sequences including a single motif), 2C (for the sense sequences including multiple motifs), or 2D (for the antisense sequences including multiple motifs). The black lines show the ROCs of the model using all DNA shape parameters, and the red lines show the ROCs of the model using only the preferentially selected parameters shown in Supplementary Figure 2. Ten lines show the result of 10 times validation pairing the positive and negative samples in LOOCV. The vertical axis of the ROC curve indicates the “Sensitivity”, which is the true positive rate, and the horizontal axis indicates the “1 - Specificity”, which is the false positive rate. *p* values were calculated using the Student’s *t* test. \**p* < 0.05.

## Discussion

This study was designed to evaluate the underlying mechanisms regulating gene expression in response to *A. oryzae* TF, AoXlnR, from the viewpoint of DNA conformation around the binding sites. We used datasets containing information on the binding sites of AoXlnR in the *A. oryzae* genome and their differential expression in response to the overexpression of this TF to construct mathematical models explaining the differential regulation of AoXlnR binding sites [6, 18].

We first determined the impact of each of the 194 parameters for each AoXlnR recognition region using 3D DNA shape information. These values were then used to construct individual prediction models, followed by applying to predict responsiveness to this TF. However, the AUCs of the models constructed using SVM were 0.644 and 0.522 for the sense and antisense strand sides, respectively, indicating that the accuracy of the constructed models was insufficient to reliably distinguish between DEG and non-DEG promoters (Fig. 2, Supplementary Fig. 1). These results suggest that the inclusion of parameters with small contribution to the differential expressions via AoXlnR prevented the construction of accurate prediction models.

Thus, we selected the top parameters in the order of their coefficient contributing to the discrimination of differential expression to reduce the number of explanatory variables used in the prediction models. These models were then reconstructed using two to ten selected parameters. These models displayed AUCs of more than 0.7 marking a significant improvement compared to the above models using all parameters (Fig. 2, Supplementary Fig. 1). It should be noted that certain DNA structural parameters were included in most of the models with higher AUCs. In fact, one of the parameters, down_17_ProT, that is determined by the sequences around 17 b downstream of the AoXlnR binding motifs, was preferentially selected. In the case of model construction using the antisense strand of the motif, the DNA shape parameters related to 15 b upstream of the binding motifs were observed at a relatively high frequency. These preferentially selected parameters were calculated from the structure of sequences outside of the conventional binding motif sequences, 5′-GGCTGA-3′ or 5′-GGCTAA-3′. This fact strongly suggests the existence of a regulatory mechanism for the expression of AoXlnR, which has not been revealed by conventional analyses based only on TF-binding motif information. We are currently planning to introduce mutations in the promoter region to alter the down_17_ProT parameters to understand how this impacts gene regulation.

Next, we further classified the MPDs by the presence of other motifs in the same promoter region, and then subjected the classified dataset to the construction of a new set of models, followed by the evaluation of the performance of resultant models. The results demonstrated that models constructed using the selected parameters which contributed to the model performance showed higher AUC than the models constructed under conditions with no classification by the number of motifs, including those using antisense strands (Figs. 3, Supplementary Fig. 2). We also randomly stratified the dataset including DNA shape parameters of MPD to validate the effect of stratification by the number of models on the accuracy of the model. Although the randomly stratified prediction models showed also high values with that of the model constructed according to the motif of the sense strands, the models based on the antisense strands of binding motifs showed higher AUC than that of the randomly stratified prediction models (Supplementary Fig. 3). Furthermore, preferentially selected parameters were different compared to the model using MPDs from the promoters including multiple motifs (Supplementary Fig. 4). For example, the models based on multiple parameters selected up_17_Roll, up_7_Roll and up_19_Roll as the highly contributed parameters, whereas the randomly stratified model selected up_19_Roll, down_20_down and down_15_MGW. These results imply that different parameters contribute to the differential expressions depending on the number of binding motifs in the promoter.

In the models based on the DNA shape parameters around the binding motifs in promoters of sense strand including only one motif, down_23_ProT was preferentially selected as in all MPD-based models (Supplementary Fig. 2A), while up_17_Roll was preferentially selected in the models produced for promoter with several binding motifs (Supplementary Fig. 2C). In contrast, the models produced using the antisense strand showed a preference for up_17_ProT or up_11_HelT where the models were constructed for single or multiple motif entries, respectively (Supplementary Figs. 2B and D). The difference in the preferred DNA shape parameters among these models resulted in improved AUCs, suggesting that the regulatory mechanism of AoXlnR in cells depends on the number of motifs in the promoter region.

Interestingly, *xynF1* (AO090103000423), which is known to be under the control of AoXlnR [21], was more accurately discriminated as a gene regulated by AoXlnR when the sequence around the MPD in the promoter was evaluated using models constructed from MPDs with multiple motifs (100%) compared to the models without classification (10%). On the other hand, randomly classified models provided around 50% of *xynF1*’s correct discrimination rate, which is much lower than that of the models using dataset of MPDs with multiple motifs (data not shown). Taken together, these results suggest that the mechanisms of expression regulation by AoXlnR in cells can be classified by the number and orientation of the AoXlnR binding motifs.

The DNA shape information used in this study was recently reported to play an important role in TF-DNA interactions [14, 22, 23]. These reports focus on the relationship between TF-DNA interactions and the DNA structure around the binding sites. Based on these approaches, we examined whether it is possible to distinguish between TF-DNA interactions observed only in *in vitro* binding reactions and TF-DNA interactions in which TF binds linked to its gene regulation in cells and are presumed to regulate the expression of downstream genes, in terms of the DNA structure around the binding sites. Our results showed that some of the structural parameters calculated from DNA sequences around the AoXlnR binding sites were closely related to the differential expression of the regulatory genes in the cell as well as in *in vitro* binding between AoXlnR and its target binding sites.

In *Aspergillus oryzae*, a paralogous TF of AoXlnR, AraR regulates similar genes involved in the catabolism pentose pathway. Although AraR and AoXlnR are similar, their target genes and target sequences are slightly different. In 2018, Ishikawa et al. reported that these TFs demonstrate no cooperative binding, but do display competitive bindings at their shared sites [24]. Therefore, the existence of AraR binding may influence the DNA conformation of AoXlnR binding sites in specific genes.

Li *et al*. reported nine additional DNA shape parameters, including four inter-bp or bp-step parameters (Rise, Shift, Slide, and Tilt) and five intra-bp or bp parameters (Buckle, Opening Shear, Stagger, and Stretch) in 2017 [25]. The construction of models for predicting the differential expression with improved accuracy may be accomplished by adding these nine parameters to the four structural parameters used in this study.

Furthermore, our models have no consideration of any other factors involved in transcriptional regulation, including chromatin accessibility, and cytosine methylation. For example, chromatin condensation forms heterochromatin and prevents TFs from binding to their preferred sequences [26, 27]. By adding the information of such epigenetic modifications as new parameters, more accurate model can be constructed for the regulatory mechanisms of various TFs, including AoXlnR.

## Materials and Methods

### Calculating the structural parameters for the regions surrounding the AoXlnR-binding motifs using DNA shape parameters

In our previous study, we divided the set of detected genes of *A. oryzae* into two groups based on their differential expression in response to over-expression of AoXlnR: 1) genes that retain one or more AoXlnR-binding sites in their promoter regions and are expression-responsive to AoXlnR and 2) genes that retain AoXlnR binding sites in the promoter regions but were not sensitive to AoXlnR overexpression [6]. Considering the binding profile of AoXlnR, in which AoXlnR preferentially binds to 5′-GGCT(G/A)A-3′ [6, 18], the number and position of 5′-GGCTGA-3′ or 5′-GGCTAA-3′ motif were obtained from 1000 bp upstream of the start codon of genes in *A.oryzae*, generating 50 b sequences that include the AoXlnR binding motif and both sides of adjacent 22 b sequences. Here we refer to the binding motifs from the promoters of genes that are responsive to overexpression as “Motif in the Promoter region of DEGs” (MPD) and motifs in the promoters of genes unresponsive to overexpression as “Motif in the Promoter region of non-DEGs” (MPnD). Note that the binding motif closest to the start codon was selected when multiple motifs were present in the promoter region.

We then calculated the DNA shape parameters (HelT, MGW, ProT, and Roll values) for each of these 50 b sequences using the DNAshapeR package from R (version 1.16.0; https://bioconductor.org/packages/release/bioc/html/DNAshapeR.html), which uses a pentamer-based prediction model built from all-atom Monte Carlo simulations of DNA structures [13, 19]. The resulting dataset contained 194 parameter values (MGW:48, ProT:48, HelT:49, and Roll:49), including the mean and standard deviation over all positions [20] for every 50 b unit around 63 MPDs (Sense strand 34 and Antisense strand 29, Table 1) and 1506 MPnDs (Supplementary Tables 1 and 2).

**Table 1.**
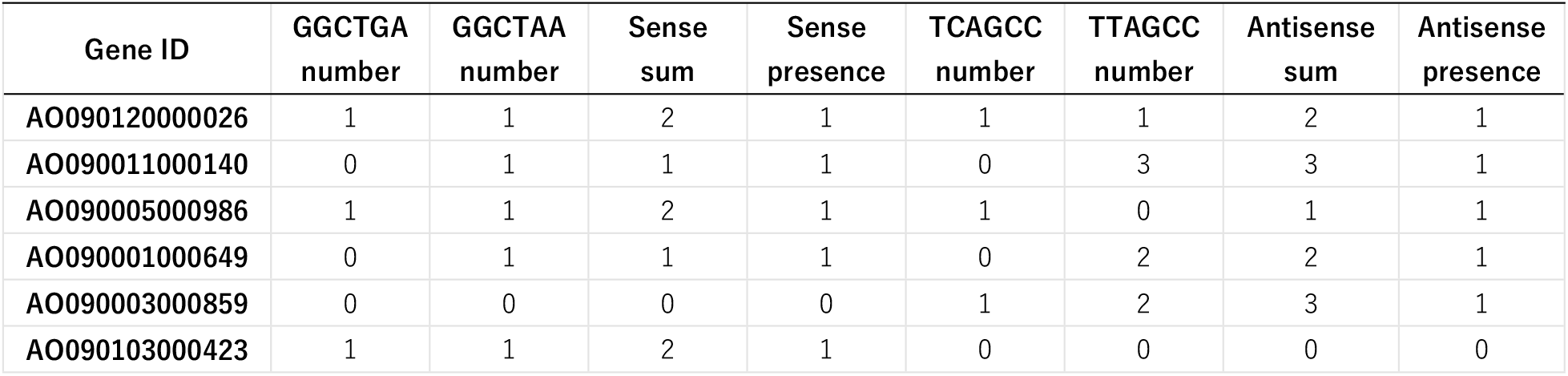
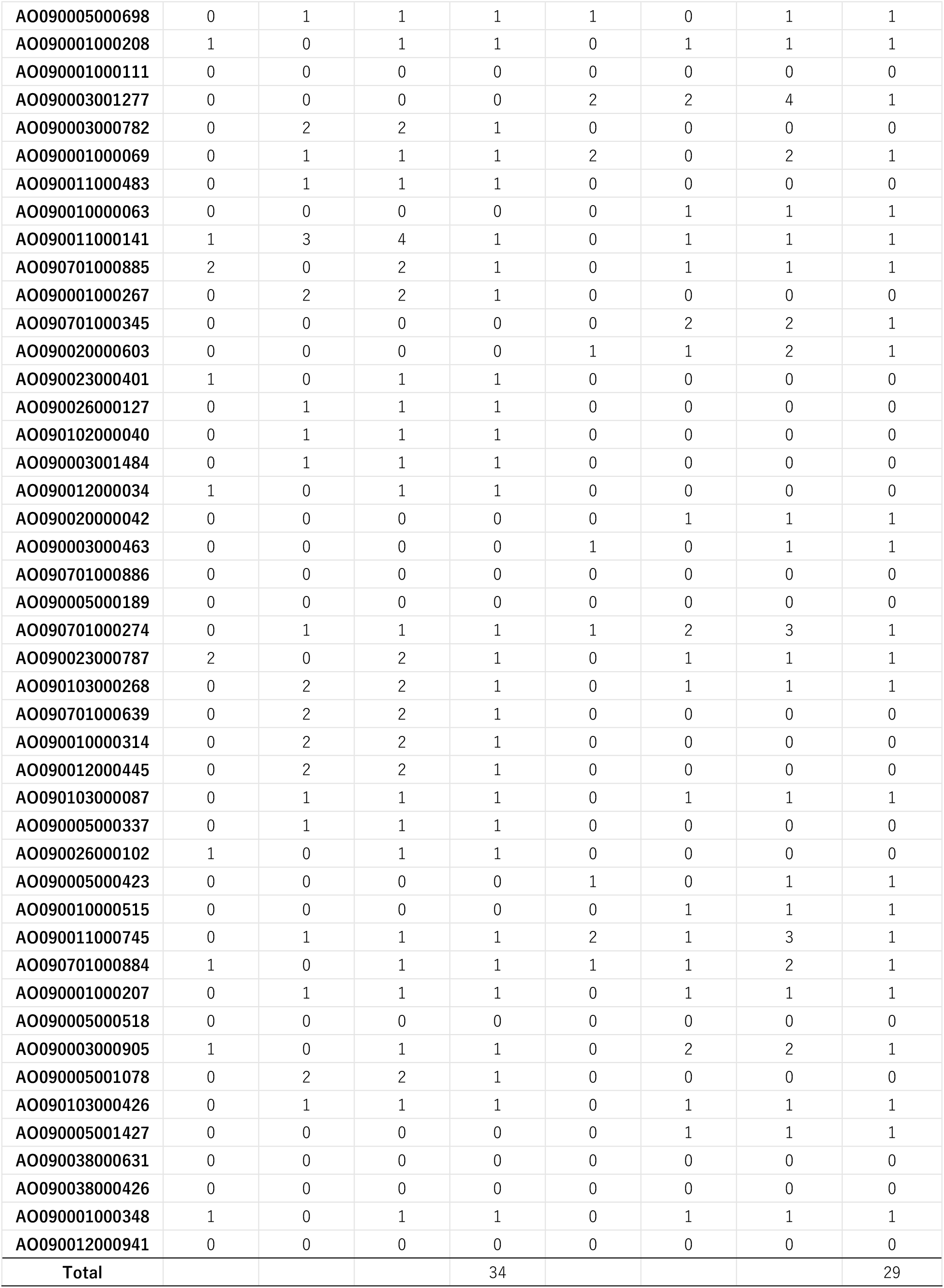
The number of AoXlnR binding motif in the promoter of MPDs.

| Gene ID | GGCTGA<br>number | GGCTAA<br>number | Sense<br>sum | Sense<br>presence | TCAGCC<br>number | TTAGCC<br>number | Antisense<br>sum | Antisense<br>presence |
| --- | --- | --- | --- | --- | --- | --- | --- | --- |
| AO090120000026 | 1 | 1 | 2 | 1 | 1 | 1 | 2 | 1 |
| AO090011000140 | 0 | 1 | 1 | 1 | 0 | 3 | 3 | 1 |
| AO090005000986 | 1 | 1 | 2 | 1 | 1 | 0 | 1 | 1 |
| AO090001000649 | 0 | 1 | 1 | 1 | 0 | 2 | 2 | 1 |
| AO090003000859 | 0 | 0 | 0 | 0 | 1 | 2 | 3 | 1 |
| AO090103000423 | 1 | 1 | 2 | 1 | 0 | 0 | 0 | 0 |
| AO090005000698 | 0 | 1 | 1 | 1 | 1 | 0 | 1 | 1 |
| AO090001000208 | 1 | 0 | 1 | 1 | 0 | 1 | 1 | 1 |
| AO090001000111 | 0 | 0 | 0 | 0 | 0 | 0 | 0 | 0 |
| AO090003001277 | 0 | 0 | 0 | 0 | 2 | 2 | 4 | 1 |
| AO090003000782 | 0 | 2 | 2 | 1 | 0 | 0 | 0 | 0 |
| AO090001000069 | 0 | 1 | 1 | 1 | 2 | 0 | 2 | 1 |
| AO090011000483 | 0 | 1 | 1 | 1 | 0 | 0 | 0 | 0 |
| AO090010000063 | 0 | 0 | 0 | 0 | 0 | 1 | 1 | 1 |
| AO090011000141 | 1 | 3 | 4 | 1 | 0 | 1 | 1 | 1 |
| AO090701000885 | 2 | 0 | 2 | 1 | 0 | 1 | 1 | 1 |
| AO090001000267 | 0 | 2 | 2 | 1 | 0 | 0 | 0 | 0 |
| AO090701000345 | 0 | 0 | 0 | 0 | 0 | 2 | 2 | 1 |
| AO090020000603 | 0 | 0 | 0 | 0 | 1 | 1 | 2 | 1 |
| AO090023000401 | 1 | 0 | 1 | 1 | 0 | 0 | 0 | 0 |
| AO090026000127 | 0 | 1 | 1 | 1 | 0 | 0 | 0 | 0 |
| AO090102000040 | 0 | 1 | 1 | 1 | 0 | 0 | 0 | 0 |
| AO090003001484 | 0 | 1 | 1 | 1 | 0 | 0 | 0 | 0 |
| AO090012000034 | 1 | 0 | 1 | 1 | 0 | 0 | 0 | 0 |
| AO090020000042 | 0 | 0 | 0 | 0 | 0 | 1 | 1 | 1 |
| AO090003000463 | 0 | 0 | 0 | 0 | 1 | 0 | 1 | 1 |
| AO090701000886 | 0 | 0 | 0 | 0 | 0 | 0 | 0 | 0 |
| AO090005000189 | 0 | 0 | 0 | 0 | 0 | 0 | 0 | 0 |
| AO090701000274 | 0 | 1 | 1 | 1 | 1 | 2 | 3 | 1 |
| AO090023000787 | 2 | 0 | 2 | 1 | 0 | 1 | 1 | 1 |
| AO090103000268 | 0 | 2 | 2 | 1 | 0 | 1 | 1 | 1 |
| AO090701000639 | 0 | 2 | 2 | 1 | 0 | 0 | 0 | 0 |
| AO090010000314 | 0 | 2 | 2 | 1 | 0 | 0 | 0 | 0 |
| AO090012000445 | 0 | 2 | 2 | 1 | 0 | 0 | 0 | 0 |
| AO090103000087 | 0 | 1 | 1 | 1 | 0 | 1 | 1 | 1 |
| AO090005000337 | 0 | 1 | 1 | 1 | 0 | 0 | 0 | 0 |
| AO090026000102 | 1 | 0 | 1 | 1 | 0 | 0 | 0 | 0 |
| AO090005000423 | 0 | 0 | 0 | 0 | 1 | 0 | 1 | 1 |
| AO090010000515 | 0 | 0 | 0 | 0 | 0 | 1 | 1 | 1 |
| AO090011000745 | 0 | 1 | 1 | 1 | 2 | 1 | 3 | 1 |
| AO090701000884 | 1 | 0 | 1 | 1 | 1 | 1 | 2 | 1 |
| AO090001000207 | 0 | 1 | 1 | 1 | 0 | 1 | 1 | 1 |
| AO090005000518 | 0 | 0 | 0 | 0 | 0 | 0 | 0 | 0 |
| AO090003000905 | 1 | 0 | 1 | 1 | 0 | 2 | 2 | 1 |
| AO090005001078 | 0 | 2 | 2 | 1 | 0 | 0 | 0 | 0 |
| AO090103000426 | 0 | 1 | 1 | 1 | 0 | 1 | 1 | 1 |
| AO090005001427 | 0 | 0 | 0 | 0 | 0 | 1 | 1 | 1 |
| AO090038000631 | 0 | 0 | 0 | 0 | 0 | 0 | 0 | 0 |
| AO090038000426 | 0 | 0 | 0 | 0 | 0 | 0 | 0 | 0 |
| AO090001000348 | 1 | 0 | 1 | 1 | 0 | 1 | 1 | 1 |
| AO090012000941 | 0 | 0 | 0 | 0 | 0 | 0 | 0 | 0 |
| Total |  |  |  | 34 |  |  |  | 29 |

### Constructing models for predicting AoXlnR-dependent differential expression using a discriminant algorithm

We constructed a set of supervised learning models to predict whether AoXlnR-binding motifs, 5′-GGCT(G/A)A-3′ in each gene promoter is in a DEG or non-DEG promoter region. The expression profiles of each of the 5′-GGCT(G/A)A-3′ motif were determined using unpublished microarray analysis data of AoXlnR overexpression strain TFX2 and AoXlnR disruption strain SK253 conducted in a previous study [18]. In this study, we used a support vector machine (e1071 package; version 1.7-3; https://cran.r-project.org/web/packages/e1071) on R (version 4.0.1; https://cran.ism.ac.jp/) as the binary classification algorithm.

To balance the difference in size between the number of MPD and MPnD, we selected the same number of MPnDs that showed the lowest response to AoXlnR overexpression as MPDs and synthesized a 50 b dataset consisting of the same number of MPDs and MPnDs. The expression variables, the DNA shape parameters were tagged with their labels; 1 for MPD, 0 for MPnD. The performance of prediction model was validated using leave-one-out cross validation. The area under the receiver operating characteristic curve (AUC) was used as the model evaluation criteria. These AUC values were calculated as the average from 10 sampling rounds.

The practical flow of building a prediction model is briefly described below (Fig. 1). First, two gene expression profiles were compared to make two categories of genes with AoXlnR-binding sites: the AoXlnR overexpression responsive genes, and AoXlnR overexpression non-responsive genes. Second, from the gene lists, the MPD and MPnD was searched. Third, from each MPD or MPnD, their sequence data of 50 b was converted into the DNA shape parameters using DNAshapeR. Fourth, using these datasets, model for predicting MPD or MPnD was constructed. In the construction of prediction model, the DNA shape parameters that significantly contribute to the differential expression of genes with the AoXlnR were ranked by calculating parameter coefficients from the SVM linear kernel, and selected parameters with a higher coefficient. Those prediction models were constructed using either (a) all DNA structure parameters, or 2 to 10 selected parameters (2 to 10 parameters) in the order of their coefficient (Supplementary Fig 1). Additionally, we constructed and evaluated a set of predictive models using datasets where MPD and MPnD data were further stratified on the number of motifs per promoter, i.e., one or more than one, using the same procedure described above. In parallel, we randomly stratified the dataset including DNA shape parameters of MPD as the same procedure with the preparation of the stratified dataset considering the number of motifs. These randomly stratified dataset was used to construct prediction models (Supplementary Fig. 2).

## Acknowledgments

We thank Tetsuo Kobayashi of Nagoya University for providing the microarray analysis data. This work was supported in part by JSPS KAKENHI Grant-in-Aid for Scientific Research(C) (Grant Number JP19K05766), JSPS KAKENHI Grant-in-Aid for JSPS fellow (Grant Number JP20J15715). This work was also financially supported in part by Mizutani Scholarship.

## Supporting information captions

**Supplementary Table 1. DNA shape parameter values of sense strand used in this study.**

**Supplementary Table 2. DNA shape parameter values of anti-sense strand used in this study.**

**Supplementary Figure 1. Parameters used and AUC values calculated in each prediction model using non-stratified dataset of DNA shape parameters.**

Weight shows the contribution for each model. Black circle shows parameters used in each model. The AUCs were calculated for models with 2 to 10 and all parameters. The AUC surrounded by the red line is the highest AUC, and the model with the ROC curve is shown in Fig. 2. (A) Models constructed based on binding motifs in sense strand. (B) Models constructed based on binding motifs in antisense strand.

**Supplementary Figure 2. Parameters used and AUC values calculated in each prediction model using stratified dataset of DNA shape parameters.**

Weight shows the contribution for each model. Black circle shows parameters used in each model. The AUCs were calculated for models with 2 to 10 and all parameters. The AUC surrounded by the red line is the highest AUC, and the model with the ROC curve is shown in Fig. 3. (A) Models constructed based on single motifs in sense strand. (B) Models constructed based on single motif in antisense strand. (C) Models constructed based on multiple motifs in sense strand. (D) Models constructed based on multiple motifs in antisense strand.

**Supplementary Figure 3. ROC curves and box plots amongst the various models constructed using randomly stratified dataset of DNA shape parameters.**

(A) ROC curve of the model based on the sense strands of MPDs from the promoters with same number of single motifs. (B) ROC curve of the model based on the antisense strands of MPDs from the promoters with same number of single motifs. (C) Box plots of AUC values obtained from ROC curve of the model based on the sense strands of MPDs with same number of single motifs. (D) Box plots of AUC values obtained from ROC curve of the model based on the antisense strands of MPDs with same number of single motifs. (E) ROC curve of the model based on the sense strands of MPDs with same number of multiple motifs. (F) ROC curve of the model based on the antisense strands of MPDs with same number of multiple motifs. (G) Box plots of AUC values obtained from ROC curve of the model based on the sense strands of MPDs with same number of multiple motifs. (H) Box plots of AUC values obtained from ROC curve of the model based on the antisense strands of MPDs with same number of multiple motifs. Each model was constructed using DNA shape parameters calculated from 50 b sequences containing AoXlnR binding motifs. Here we displayed the ROC curves of the models with the highest AUC for discriminant model using stratified MPDs selected parameters shown in Supplementary Figure 4A (for the sense sequences including a single motif), 4B (for the antisense sequences including a single motif), 4C (for the sense sequences including multiple motifs), or 4D (for the antisense sequences including multiple motifs). The black lines show the ROCs of the model using all DNA shape parameters, and the red lines show the ROCs of the model using only the preferentially selected parameters shown in Supplementary Figure 4. Ten lines show the result of 10 times validation pairing the positive and negative samples in LOOCV. The vertical axis of the ROC curve indicates the “Sensitivity”, which is the true positive rate, and the horizontal axis indicates the “1 - Specificity”, which is the false positive rate. *p* values were calculated using the Student’s *t* test. \**p* < 0.05.

**Supplementary Figure 4. Parameters used and AUC values calculated in each prediction model using randomly stratified dataset of DNA shape parameters.**

Weight shows the contribution for each model. Black circle shows parameters used in each model. The AUCs were calculated for models with 2 to 10 and all parameters. The AUC surrounded by the red line is the highest AUC, and the model with the ROC curve is shown in the Supplementary Figure 3. (A) Models constructed based on single motifs in sense strand. (B) Models constructed based on single motif in antisense strand. (C) Models constructed based on multiple motifs in sense strand. (D) Models constructed based on multiple motifs in antisense strand.

